# The Cost of Fame: Strong Biases in Comparative Oncology of Captive Species

**DOI:** 10.1101/2025.09.25.675237

**Authors:** Antoine M. Dujon, Jonas Courtalon, Klara Asselin, Frédéric Thomas

## Abstract

Comparative oncology is a rapidly expanding field that seeks to explain variation in cancer risk across species by examining trends between tumour prevalence and key risk factors such as body mass, longevity, life history traits, and mutation rates. These trends are then used to address fundamental questions in the field, including the discovery of potential novel anti-cancer therapies, improvements to species conservation efforts, and understanding how cancer has influenced the evolution of multicellularity. They thus must be robust. This study demonstrates that when estimated on captive species those trends are heavily influenced by their scientific and public popularity, and that accounting for this bias can substantially alter their direction and magnitude. Hence, we reanalysed published captive vertebrate datasets examining the associations between neoplasia, malignancy, and lethal tumour prevalences with body mass, longevity, life history traits, and germinal cells mutation rates. When we included proxies of species popularity in our analyses, the previously reported weak effect of body mass on tumour and malignancy prevalences disappeared entirely. Similarly, the previously reported positive relationship between germline mutation rate and cancer mortality was eliminated after controlling for popularity bias. For life history traits, the effect of clutch size on cancer neoplasia and malignancy prevalences in birds doubled in magnitude, and while the negative trend between gestation length and tumour prevalence in mammals was not greatly affected, our analyses revealed that baseline tumour prevalences were underestimated for popular animals. Finally, the previously observed association between hemochorial placentation and cancer mortality in mammals was eliminated when confounding variables were included. Collectively, these results demonstrate that current comparative analyses based on tumour prevalence in captive animals are heavily influenced by species’ scientific and public popularities. Future studies utilising such datasets should incorporate measures of species popularity as confounding variables to ensure more robust conclusions and misleading research directions.

## Introduction

Over the past decade, comparative oncology has gained momentum as a field to explore the evolutionary foundations of cancer resistance (e.g. Boddy et al. 2020a; Dujon et al. 2023a, b, 2025; Kapsetaki et al. 2024a, 2025; Matthews et al. 2025; Møller et al. 2017; Vincze et al. 2022). From whales to elephants to naked mole rats, researchers have searched for patterns explaining why some species appear less susceptible to cancer than others (Manuel Vazquez *et al*. 2022; Nunney 2022). These studies have recently fed into broader and sometimes heated discussions on the role of tumour suppression in vertebrates challenging our assumptions about cancer risk in large, long-lived animals (Butler *et al*. 2025; Compton *et al*. 2025; Dujon *et al*. 2025; Lynch 2025; Simons 2025). The arguments presented in these debates typically rely on information obtained from comparative analyses of cancer prevalence in captive or wild mammals to address fundamental questions about cancer ecology and evolution (Dujon *et al*. 2021). It is therefore crucial that the trends observed in these studies, which scientists use to build their interpretations, are robust, given the risk of false directions and dead ends. The implications are significant, as comparative oncology studies aim to facilitate the potential discovery of novel anti-cancer therapies (Boddy *et al*. 2020b), improve species conservation efforts (Dujon *et al*. 2024b; Hamede *et al*. 2020), and more fundamentally, understand how cancer has shaped the evolution of multicellularity (Aktipis *et al*. 2015; Albuquerque *et al*. 2018; Boutry *et al*. 2020; Dujon *et al*. 2022; Jacqueline *et al*. 2017; Thomas *et al*. 2017).

However, amid this flourishing literature, one essential issue has been surprisingly overlooked: the role of sampling effort bias. Sampling bias is well documented in modern science (Ducatez & Lefebvre 2014; Hickisch *et al*. 2019; Hughes *et al*. 2021; Tam *et al*. 2022). As a result, certain domains of comparative biology, such as parasite ecology, routinely account for variation in research attention (Kamiya *et al*. 2014; Poulin *et al*. 2023). However, comparative oncology analyses in captive mammals have largely proceeded under the assumption that tumour detection probability depends solely on the number of necropsies performed for a given species and potential risk factors such as life history traits, diet, or parasite exposure (Butler *et al*. 2025; Compton *et al*. 2024; Dujon *et al*. 2023a, b, 2025; Kapsetaki *et al*. 2024a, 2025; Vincze *et al*. 2022). This approach fails to consider that prevalence estimates may be significantly affected by our varying knowledge of and interest in different species, which can influence the likelihood that a tumour is correctly detected and diagnosed in a deceased animal. This is particularly critical given that cancer, like parasitism (e.g. Poulin et al. 2023a), often requires targeted investigation to be detected, especially outside of model species (i.e. the pathologist must know what to look for). For example, in human medicine, it is documented that wealthier populations, with better access to screening and healthcare, are more likely to be diagnosed with cancer, even when this increased detection does not translate into higher mortality (Welch & Fisher 2017). Most current comparative oncology studies acknowledge potential sampling biases due to detection probability in their discussions but do not include them in their statistical analyses, with a small number of studies such as Butler et al. (2025) clearly lacking of cautiousness (see the discussions of Compton et al. 2025, Dujon et al. 2025 and Simons 2025 in that regard). There is therefore a critical need for proper quantification of how these biases may affect the conclusions drawn from comparative oncology studies.

In this study, we show that those diagnosis and sampling biases do exist in comparative oncology, and that failing to account for them can lead to fundamentally misleading conclusions. Revisiting several recent studies testing different hypotheses that may explain tumour prevalence in captive mammals (Compton *et al*. 2024; Dujon *et al*. 2023b; Kapsetaki *et al*. 2024a; Vincze *et al*. 2022), we reproduced their original analyses and then introduced simple yet critical proxies for sampling effort aiming to quantify the scientific and general public interests in each species. Strikingly, once these variables are included, previously reported significant associations involving tumour prevalence, body mass or germinal cell mutation rate vanish, while patterns involving reproductive traits strengthen. These findings suggest that much of what has been inferred about cancer susceptibility across species may have been shaped, in part, by research visibility rather than biological reality. Addressing this blind spot is essential if we are to build a truly evolutionary understanding of cancer defences.

## Material and methods

To investigate the effect of confounding factors on comparative analysis of tumour prevalence in vertebrates we reanalysed datasets originally published by various teams which were aimed to test some important hypotheses in the field: Peto’s paradox, the effect of germinal cell mutation rates, and the effect of reproductive traits. We focused here on three different types of tumour prevalences obtained from necropsied captive animals and verified by pathologists: neoplasia, malignancy, and lethal (i.e. when the tumour is considered to have significantly contributed to the death of the animal). We refer the reader to the original publication for the detailed information on how those datasets were collected. When needed we also describe how we complemented those datasets.

### Peto’s paradox

Peto’s paradox remains a prominent topic in cancer biology, generating extensive debate (Caulin & Maley 2011; Dujon *et al*. 2025; Nagy *et al*. 2007; Noble *et al*. 2015; Roche *et al*. 2012; Tollis *et al*. 2017). The paradox stems from the multistage carcinogenesis model, which incorporates body mass, longevity, number of cells, and cell divisions to accurately predict cancer risk within species such as humans and dogs (Albanes & Winick 1988; Nunney 2018, 2024). According to the model, larger, longer-lived species should experience an increase in cancer incidence with a strong interaction effect between those two traits (Nunney 1999). However, when measured cancer risk is compared across multiple species, the incidence is far lower than their body mass and longevity would predict using the multistage carcinogenesis model (Compton *et al*. 2024; Dujon *et al*. 2025; Vincze *et al*. 2022). This discrepancy, from which the paradox comes from, has been interpreted as evidence that species have evolved potent anti-cancer defences to counteract the risks associated with large body size and extended lifespan (Nery *et al*. 2022; Seluanov *et al*. 2018; Trivedi *et al*. 2023).

Recently, Butler *et al*. (2025) reanalysed Compton *et al*. (2024) vertebrates’ dataset and reported a small positive trend between body size and cancer risk but found no association with longevity, a pattern previously observed for both neoplasia and malignancy in the original dataset (Compton *et al*. 2025). Butler *et al*. (2025) argued that these results invalidate Peto’s paradox, a claim that has been immediately challenged, among other things because no trend with longevity was observed (Compton *et al*. 2025; Dujon *et al*. 2025; Simons 2025), because the effect size relative to the body size is small and because the positive association with cancer risk and body mass becomes non-significant when considering interaction with longevity (Dujon *et al*. 2025). In the case of cancer mortality, Vincze *et al*. (2022) found no relationship between body mass and mortality risk when restricting analyses to species in which cancer was detected, while more recently Dujon *et al*. (2025) identified a small negative trend between cancer mortality risk and body mass when reanalysing Vincze’s dataset to include both species for which cancer was and was not observed. However, this trend disappeared when body mass was considered with an interaction with longevity. The current level of evidence thus suggest that Peto’s paradox is real, but it also assumed the estimated trends are robust. A key benefit of accurately quantifying trends (or lack thereof) between body mass, longevity, and cancer risk is the potential to identify species with substantial differences between observed and predicted cancer prevalence. This approach could reveal how large species might trade off cancer protection mechanisms against energy allocated to reproduction to preserve their fitness (Boddy *et al*. 2015; Dujon *et al*. 2024a; Jacqueline *et al*. 2017). Considering the ongoing debates around Peto’s paradox we wanted to assess whether the above trends persisted after accounting for proxies of both scientific and public interest, and we thus reanalysed neoplasia, malignancy data from Compton et al. (2024), and mortality data from Vincze *et al*. (2022).

### Effect of germinal mutation rates on cancer risk

Based on a comparative analysis of 37 vertebrate species, Kapsetaki *et al*. (2024b) reported a small positive association between germline cells mutation rates and cancer mortality. However, as the authors acknowledge, the biological link between these two traits remains unclear. One hypothesis they highlight is that a higher germline mutation rate could potentially contribute to a greater hereditary cancer burden. To explore this further, we re-analysed their dataset to test whether the observed trend persisted after accounting for proxies of both scientific and public interest.

### Effect of reproductive traits on cancer risk

Several theoretical publications predict that trade-offs should exist between reproduction and anti-cancer defences, and comparative studies have been conducted to explore these potential relationships (Boddy *et al*. 2015; Dujon *et al*. 2022; Jacqueline *et al*. 2017). For example, in mammals, placentation invasiveness has been linked to cancer mortality, with mammals that exhibit intermediate placentation invasiveness experiencing higher cancer mortality risk compared to those with high placentation invasiveness. In addition, longer gestation lengths are also associated with lower mortality risk. This suggests that adaptations have evolved to counteract placental aggressiveness (Dujon *et al*. 2023b). Across vertebrate species Compton *et al*. observed a negative association between neoplasia, malignancy, and gestation length (Compton *et al*. 2024). In birds, Kapsetaki *et al*. (2024a) observed a positive relationship between neoplasia and malignancy and clutch size, but not incubation length. These promising results suggest that trade-offs between reproductive traits and cancer defences may exist in vertebrates. We therefore sought to investigate whether these trends persist when scientific and public interest are controlled for.

### Quantification of the scientific research interest

We hypothesise that tumour prevalence could be positively associated with the amount of scientific research a species has received, as greater biological knowledge may increase the likelihood of tumour detection (as observed, for example in parasites, see Poulin et al. 2023). To quantify research attention, we collected data for each species on three key metrics (Tam *et al*. 2022): (1) the total number of publications, (2) the total number of citations those publications received, and (3) the h-index, defined as the highest number *h* such that *h* publications have each been cited at least *h* times (Hirsch 2005). These metrics were obtained using the Scopus database (https://www.scopus.com) by searching for each species’ binomial nomenclature within article titles, abstracts, and keywords and calculating the metrics using the “Citation overview” tool. This approach serves as a proxy for the level of expert attention specifically directed at each species. For species that have undergone taxonomic revisions, we conducted separate searches using both the current and former names, then merged the results while carefully removing duplicates to ensure accurate counts.

### Quantification of the impact of country economics

We hypothesise that wealthier countries may invest more in conservation, zoological institutions, or advanced diagnostic methods, which could result in higher reported prevalences. This is based on the observation that conservation spending is generally correlated with a country’s per capita income (McClanahan & Rankin 2016; McKinney 2002). To investigate this effect, and since a species can be present in multiple institutions across multiple countries, we calculated a weighted mean and median Gross Domestic Product (GDP) per capita associated with each species by averaging the GDP per capita of each country, weighted by the number of institutions holding the species in that country (thereby giving more weight to countries having the species in multiple institutions). To ensure appropriate GDP time frame selection, we calculated the mean and median species GDP per capita over 5-year, 10-year, and 20-year periods (ending in 2025 and going backward). The raw GDP data time series were obtained from the International Monetary Fund website (https://www.imf.org/).

### Quantification of the general public interest

Species that attract greater public attention may be more frequently diagnosed with neoplasia or malignancies because they may face more intense public scrutiny, prompting zoo staff to thoroughly investigate causes of death for publicly valued animals (Clauss et al. 2025; Hutchins 2006). Multiple studies found that Wikipedia page views are an effective proxy for measuring public interest for a given species at large scales (Mittermeier *et al*. 2021; Roll *et al*. 2016). We thus obtained the total number of Wikipedia page views for each species by searching for their scientific names across pages in 173 different languages over the past 20 years using the Wikimedia website (https://commons.wikimedia.org/). We collected data covering the period from January 1, 2005, to January 1, 2025.

### Phylogenetic regression models

Because the collected variables often had a skewed distribution with a long tail, we log 10 transformed them when required (this included body mass, maximum longevity, gestation length, lactation length, H-index, number of institutions, number of Wikipedia views). For variables that can have a value of zero (e.g. H-index), we added 1 to all values before log 10 transforming them. Exploration of the correlation matrixes calculated from the potential confounding variables revealed significant collinearity among several of them, preventing their simultaneous inclusion in phylogenetic regression models (**Supplementary Figures 1-3**). Upon reviewing the data, we selected a representative variable for each group of highly correlated predictors. As it can be expected, a species’ H-index was strongly correlated with both its number of publications and citations, being a composite of the two, so we retained the H-index as a proxy for research interest. All GDP per capita-based metrics were also highly correlated; we therefore arbitrarily selected the mean 5-year GDP per capita for modelling and considering the very high collinearity we assumed similar effects across metrics. Likewise, the number of animals in captivity was strongly correlated with both the number of holding institutions and countries, so we retained the number of institutions as a measure of geographic distribution in captivity. Due to the strong correlation between Wikipedia views and H-index values across species, we calculated residuals from a regression of Wikipedia views against H-index for each species to address this collinearity. These residuals represent the deviation from the expected number of Wikipedia views based on a species’ H-index. For any given H-index value, a positive residual indicates that a species receives more Wikipedia views than average, while a negative residual indicates fewer views than average.

To compute the variance-covariance matrices used in our models to account for the lack of evolutionary independence between species, we used a phylogenetic tree obtained from TimeTree (www.timetree.org). Previous studies have shown that an Ornstein–Uhlenbeck process provides a good fit for modelling the evolution of cancer risk across phylogenies (Compton *et al*. 2024; Dujon *et al*. 2025; Kapsetaki *et al*. 2024a, b). Accordingly, we used the Ornstein–Uhlenbeck model to calculate the variance-covariance matrix used to structure the residuals in our models.

### Purposeful selection of variables

To investigate the effect of potential confounding variables on our models, we adapted the Purposeful Selection of Variables model selection method, which is commonly used in epidemiology to identify confounding variables in binomial regression models to the Bayesian framework (see Dujon et al. 2025 for full details on that framework). This method was selected because it is specifically designed to identify confounding variables in binomial regression models (Bursac *et al*. 2008; Hosmer Jr. *et al*. 2013). The selection procedure was conducted on species with a complete case (i.e. no missing values across the different potential confounding variables and risk factors).

The purposeful selection method was applied in four steps for each of the three tumour types (neoplasia, malignancy, lethal): (1) First, we fitted a univariable binomial regression model between tumour prevalence and each of the potential risk factors and potential confounding variables. We retained variables with a p-direction >0.90 for the next step. The p-direction is the probability that a parameter has a consistent directional effect (either consistently positive or consistently negative) based on the proportion of posterior samples that fall on the same side of zero. More traditional levels such as p-direction >0.95 can fail to identify variables known to be important (Bursac *et al*. 2008). (2) We then built a model including all the variables retained in step (1) and iteratively refitted this model by dropping each variable one at a time and investigating the effect of the dropping on the estimates of the other variables. A change in a parameter estimate above a specified level indicates that the excluded variable was important in the sense of providing a needed adjustment for one or more of the variables remaining in the model. Here we used a 25% change as a threshold, which is more conservative than the 20% recommended by Hosmer Jr (2013). It is not unreasonable to consider that the inclusion of a variable in a model that has this effect on other parameters should be included as a confounding variable in the model (3) Once this process was completed, we reintroduced the variables excluded in step (1) to ensure they did not affect the estimates of the final model (using a change in estimate of 25% as criteria). (4) Finally, we final refitted the model to interpret them with their 95% credible intervals. This process does not necessarily result in the most parsimonious model but is designed to identify and retain confounding variables that considerably affect the slopes of other parameters included in the model. This selection of variables allows to compare crude models, which include the key risk factors mentioned in the different publications above, to adjusted models and to compare their effect size, for example their odds ratio.

Following completion of the model selection procedure, we computed marginal effects to examine how confounding variables influenced tumour prevalence (Mize *et al*. 2019). These marginal effects were derived by maintaining all parameters at constant values except for the variable under investigation (held at median values, representing animals of median scientific and general public popularity). This approach enabled us to visualise the predicted changes in average tumour prevalence associated with each potential risk factor across species while controlling for confounding variables (Mize *et al*. 2019).

All Bayesian analyses in this study were conducted using the brms R package (Bürkner 2017). All models were fitted using four chains, each run for a minimum of 5,000 iterations with a 2,000-iteration burn-in period and thinning interval of 5. All models employed default uninformative priors. Model convergence was assessed through multiple diagnostic criteria: autocorrelation within chains was confirmed to be low (R < 0.1), Rhat values for all estimated parameters equalled 1, and both bulk and tail effective sample sizes exceeded 1,000. We also visually examined posterior distributions to verify they were unimodal and conducted posterior predictive checks to identify poorly fitting models (Bürkner 2017; Conn *et al*. 2018). For successfully converged models, we converted slope coefficients to odds ratios (ORs) with corresponding 95% credible intervals to facilitate interpretation (Szumilas 2010).

The code and raw datasets using in this publication are available on GitHub (https://github.com/adujon/ComparativeOncologyBiases).

## Results

### Univariate models

Species that are more extensively studied (indicated by high H-index values), widely distributed across multiple zoos, maintained in large captive populations, or possess significant public appeal and Wikipedia readership show elevated rates of neoplasia, malignancy, and mortality (**Figure 1**). In contrast, when species are housed in countries with a high GDP, the occurrence of neoplasia and malignancy decreases across vertebrate groups, though mortality risk increases among mammals (**Figure 1, 2**). These patterns indicate that such factors act as significant confounding variables and must be included as covariates when conducting comparative research on cancer prevalence in zoo animals.

**Figure 1:**
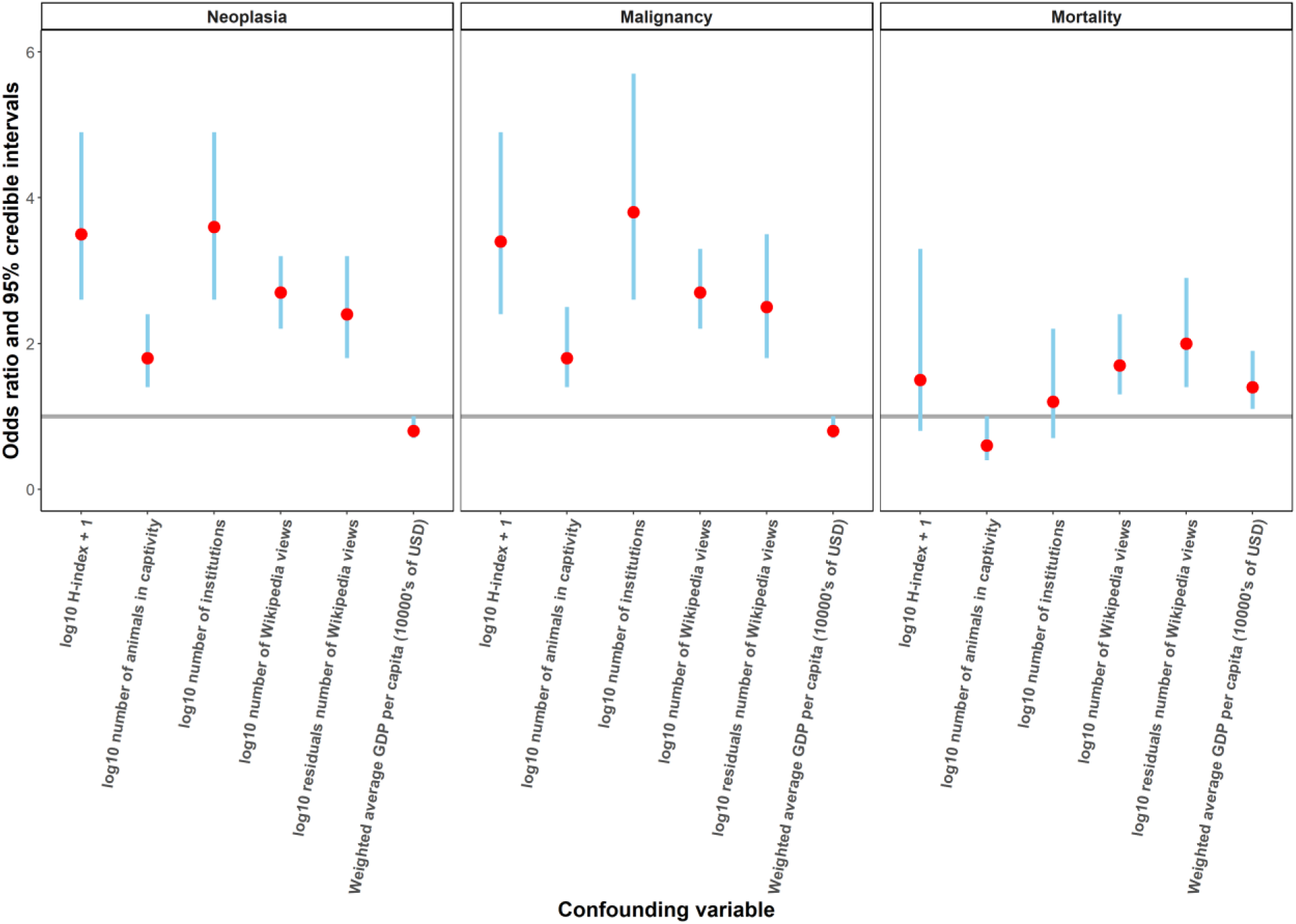
Odds ratios and associated 95% confidence intervals calculated when considering univariate models between neoplasia (n = 325 vertebrate species), malignancy (n = 325 vertebrate species) and mortality prevalences (n = 172 mammal species) and potential confounding variables in captive animals. Scatterplot and regression lines for each variable are provided in **Supplementary Figures 4-6**.

**Figure 2:**
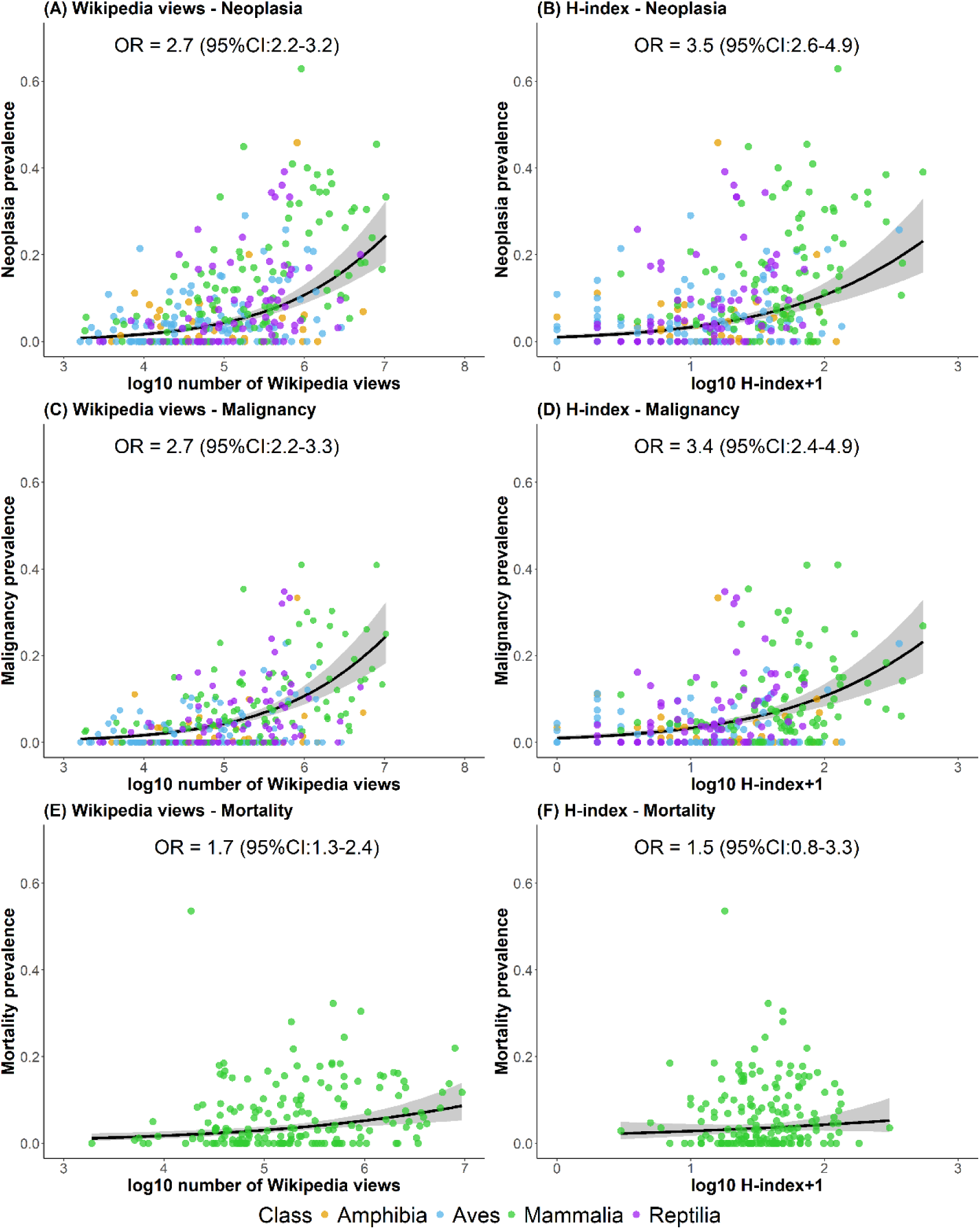
Two confounding variables showing how. (a-b) neoplasia, (c-d) malignancy and (e-f) lethal tumours prevalences are associated with the number of Wikipedia page views (a proxy of public interest) and the H-index (a proxy of scientific interest) in captive vertebrate species. The regression lines and the associated 95% credible intervals bands, along with their odds ratios, were calculated using univariate binomial phylogenetic regression models examining the relationship between species popularity metrics and tumour prevalence in captive vertebrates. Scatterplot and regression lines for the other explored metrics are presented in **Supplementary Figures 4-6**.

### Peto’s paradox

A species’ H-index, number of institutions, and the residuals of Wikipedia page views are confounding variables that influenced how longevity and body mass relate to the prevalences of neoplasia, malignancy, and lethal tumours (n = 212 species for neoplasia and malignant tumours, n = 164 species for lethal tumours, **Figure 3**).

**Figure 3:**
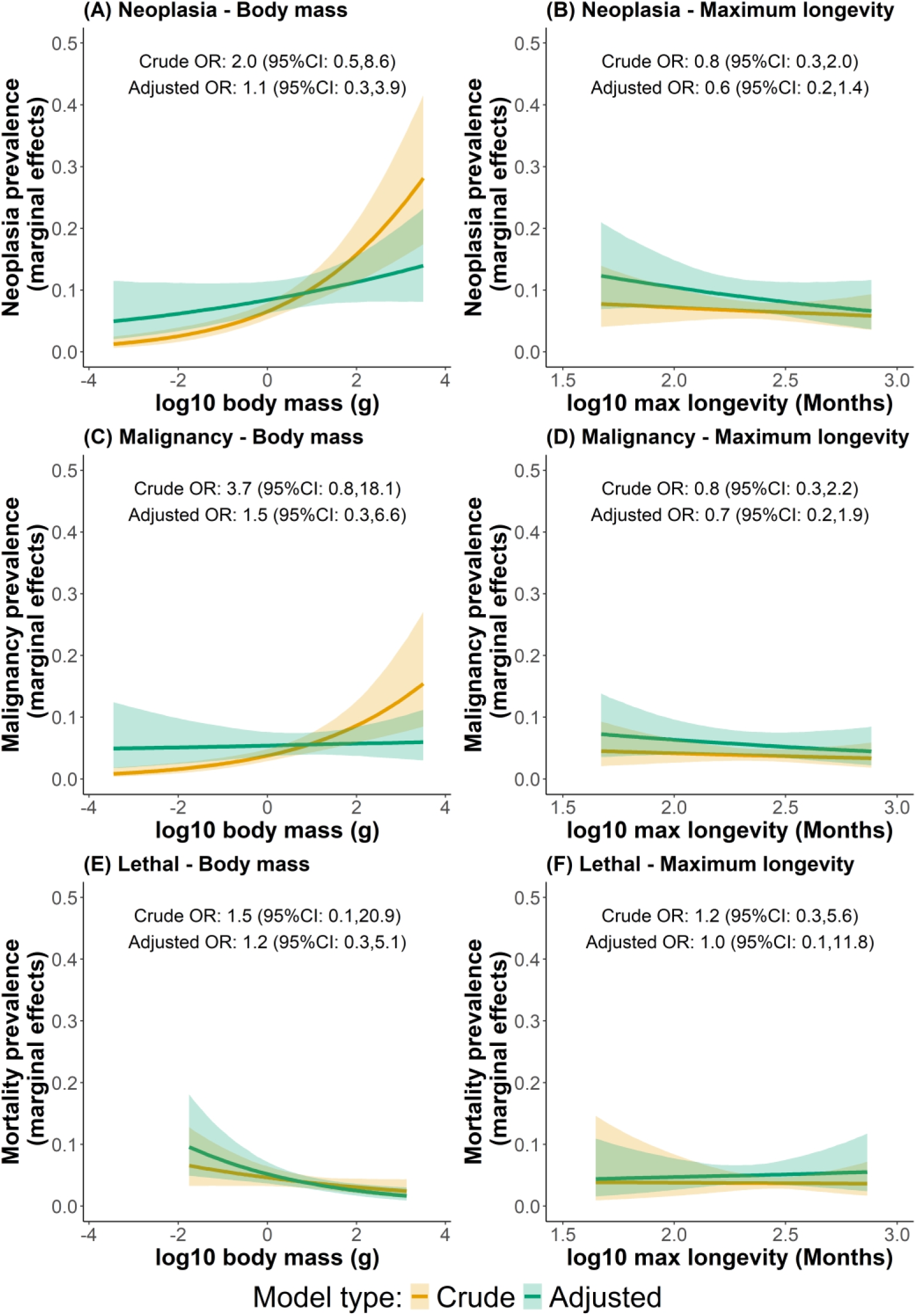
Effect of (a,c,e) body mass of vertebrates and (b,d,f) maximum longevity on the prevalence of neoplasia, malignancy and lethal tumours in vertebrates with (adjusted model, in green) and without (crude model, in yellow) accounting for confounding variables in the models. The marginal effects are shown for vertebrate species of median scientific and general public popularity.

When confounding variables are incorporated into the analysis, the already weak and non-significant relationship between body mass and the prevalence of the three tumour types becomes further attenuated when considered simultaneously with maximum longevity, a pre-requisite condition for testing Peto’s paradox. Indeed, odds ratios are reduced by half for neoplasia and malignancy, and by 25% for mortality, and none of these associations achieve statistical significance (**Figure 3a,c,d**). In comparison, the relationship between longevity and tumour prevalence remains largely unchanged when confounding variables are added to the models. While overall baseline prevalence estimates for an animal of median popularity increase slightly with the inclusion of these factors, the associations between longevity and neoplasia, malignancy, and lethal tumours remain non-significant across the full spectrum of maximum longevity values. Thus, the inclusion of confounding variables provides support to Peto’s paradox by removing the previously observed weak trend between body mass and tumour prevalence while maintaining the lack of significance for longevity associations (**Figure 3b,e,f**).

### Effect of germinal cell mutation rate on lethal tumours

The H-index of species, the weighted mean GDP of countries housing them, and the residuals of their Wikipedia page views act as confounding variables in the relationship between germline mutation rates and mortality prevalence in captive animals (n = 32 species). When these confounding variables are incorporated into the model, the odds ratio between mutation rate and mortality prevalence is reduced by close to 75% indicating minimal evidence for an association between germline mutation rates and cancer mortality in captive animals when the biases are accounted for (**Figure 4**).

**Figure 4:**
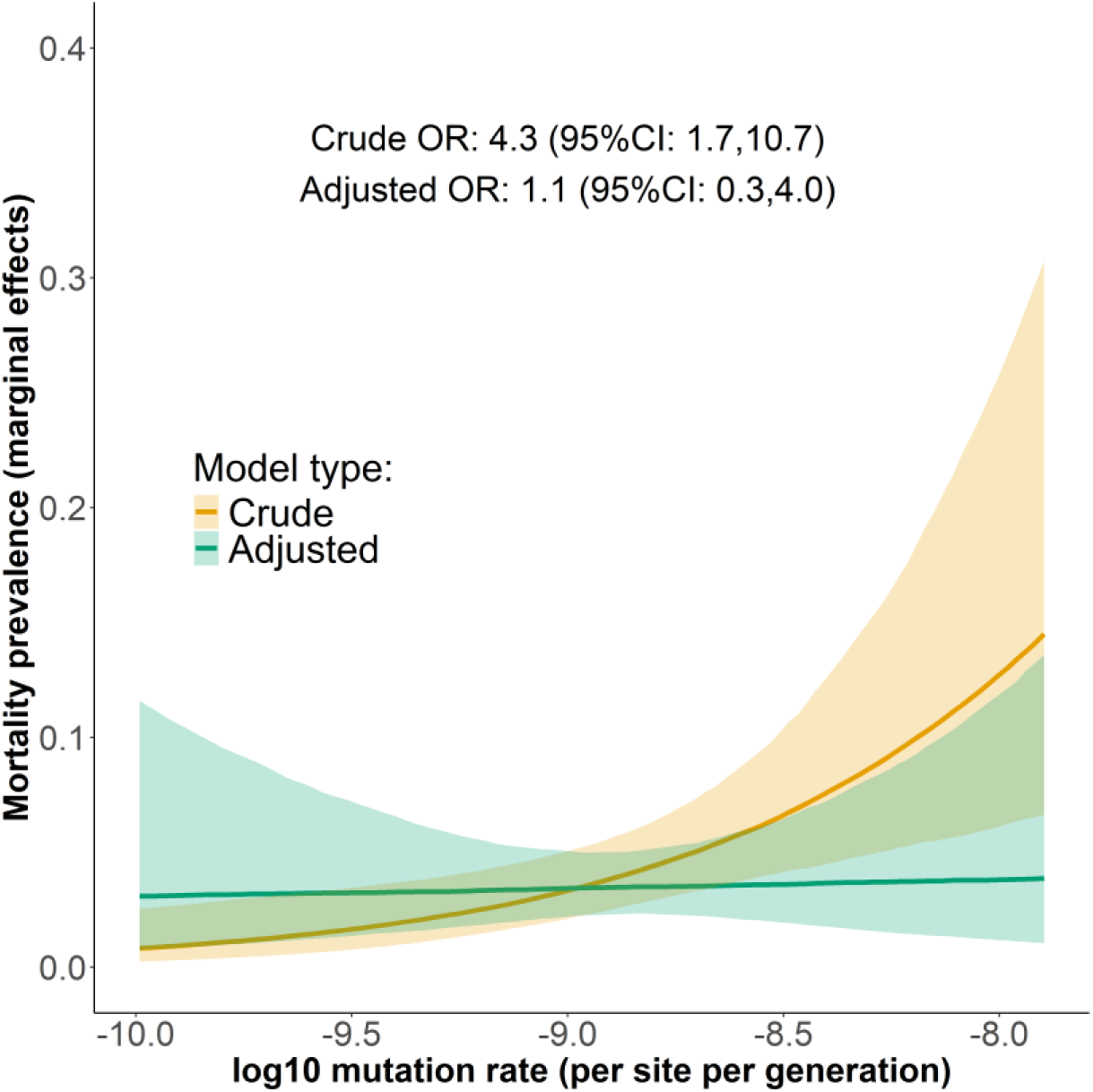
Comparison of the prevalence of lethal tumours in vertebrates (birds, reptiles, fish) in function of their germinal cell mutation rate before and after accounting for confounding variables.

### Effect of reproductive traits on cancer risk

The H-index of a species, number of institutions, weighted mean GDP, residuals of Wikipedia page views and the diet act as confounding factors for placentation type and gestation length in mammals (n = 88 species for neoplasia and malignancy tumours, n = 119 for lethal tumours). Lactation length was not retained as a risk factor in the final model. For neoplasia, including the confounding variables increases the predicted effect of placentation, but this increase is not statistically significant and is associated with considerable uncertainty (**Figure5a, c**). Similarly for malignancy, including the confounding variables does not greatly change the predicted effect of epitheliochorial and hemochorial placentation on prevalence but increases their uncertainty. The confounding effect is most notable for lethal tumour prevalence, where the effect of endotheliochorial placentation, the most invasive placentation type, is more than halved to reach the effects of the other placentation types when confounding factors are accounted for, indicating that the higher prevalences observed for this placentation type were biased (**Figure 5e**).

**Figure 5:**
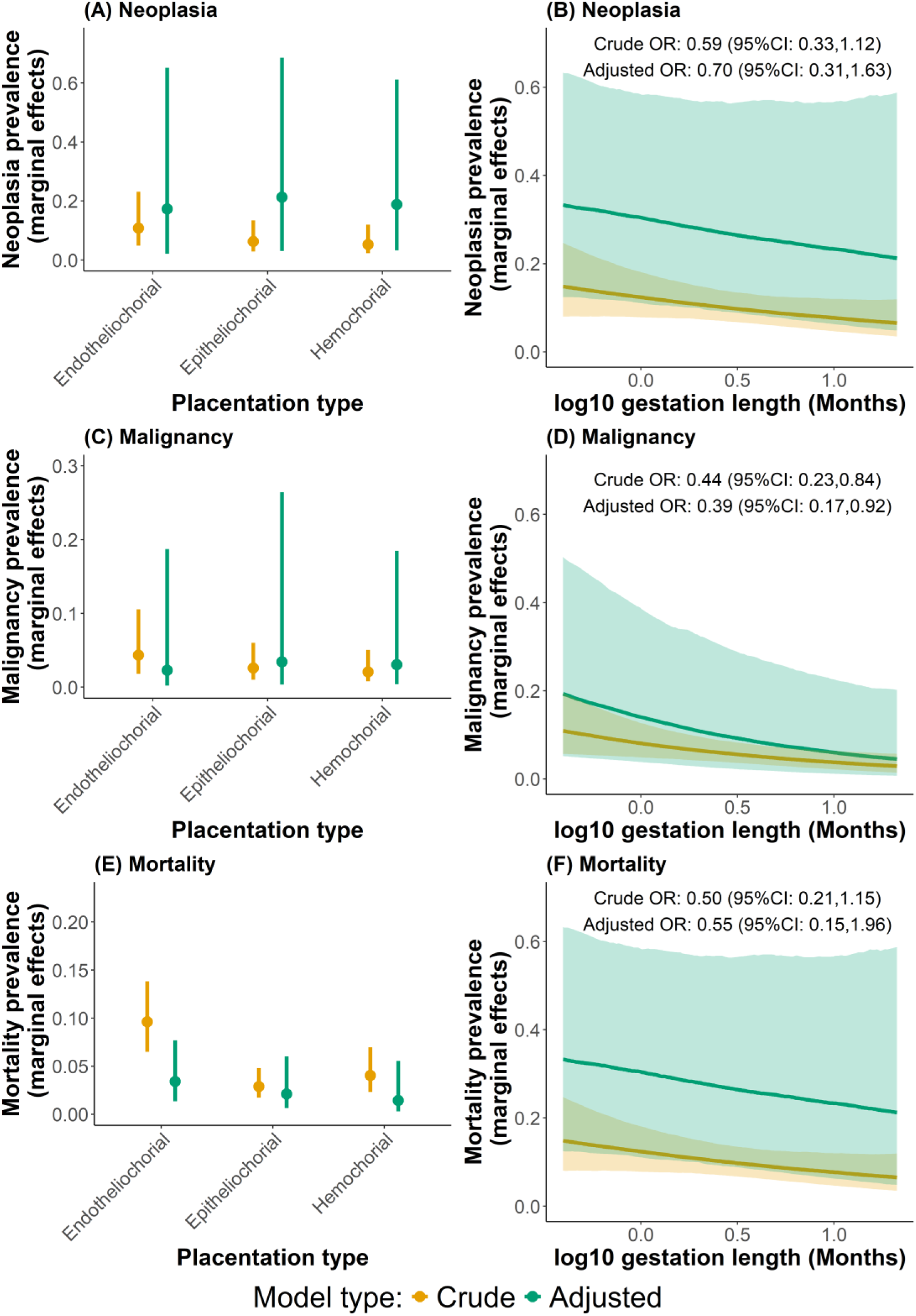
Effect of (a,c,e) placentation type and (b,d,f) gestation length on the prevalence of neoplasia, malignancy, and mortality in mammals with (adjusted model in green) and without (crude model in yellow) accounting for confounding variables in the models. The marginal effects are shown for mammalian species of median scientific and general public popularity.

While the slopes between gestation length and neoplasia, malignancy, and lethal tumour prevalence remained relatively similar, the inclusion of confounding variables substantially increased the uncertainty around these relationships. Among the three associations, only the relationship between malignancy prevalence and gestation length remained statistically significant. It also revealed an overall underestimated prevalence across the gestation length range that was masked by the confounders, with higher predicted prevalence for each of the three prevalence types (**Figure 5b,d,f**). For an animal of median scientific and public popularity this translates into a doubling in the estimated prevalence of the three tumour types.

The weighted mean GDP per capita, the H-index, and residuals of Wikipedia views act as confounding variables affecting the relationship between clutch size and neoplasia and malignancy prevalences in birds (n = 54 species). These confounders increase the odds ratios by approximately 5-6%, an effect that must be interpreted within the range of clutch size variation (1 to 24 eggs). For species with the largest clutch sizes and of median scientific and general public popularity, inclusion of these confounding variables results in a doubling of estimated neoplasia and malignancy prevalences attributable to clutch size (**Figure 6**). Thus, the effect of reproduction in those birds was previously underestimated.

**Figure 6:**
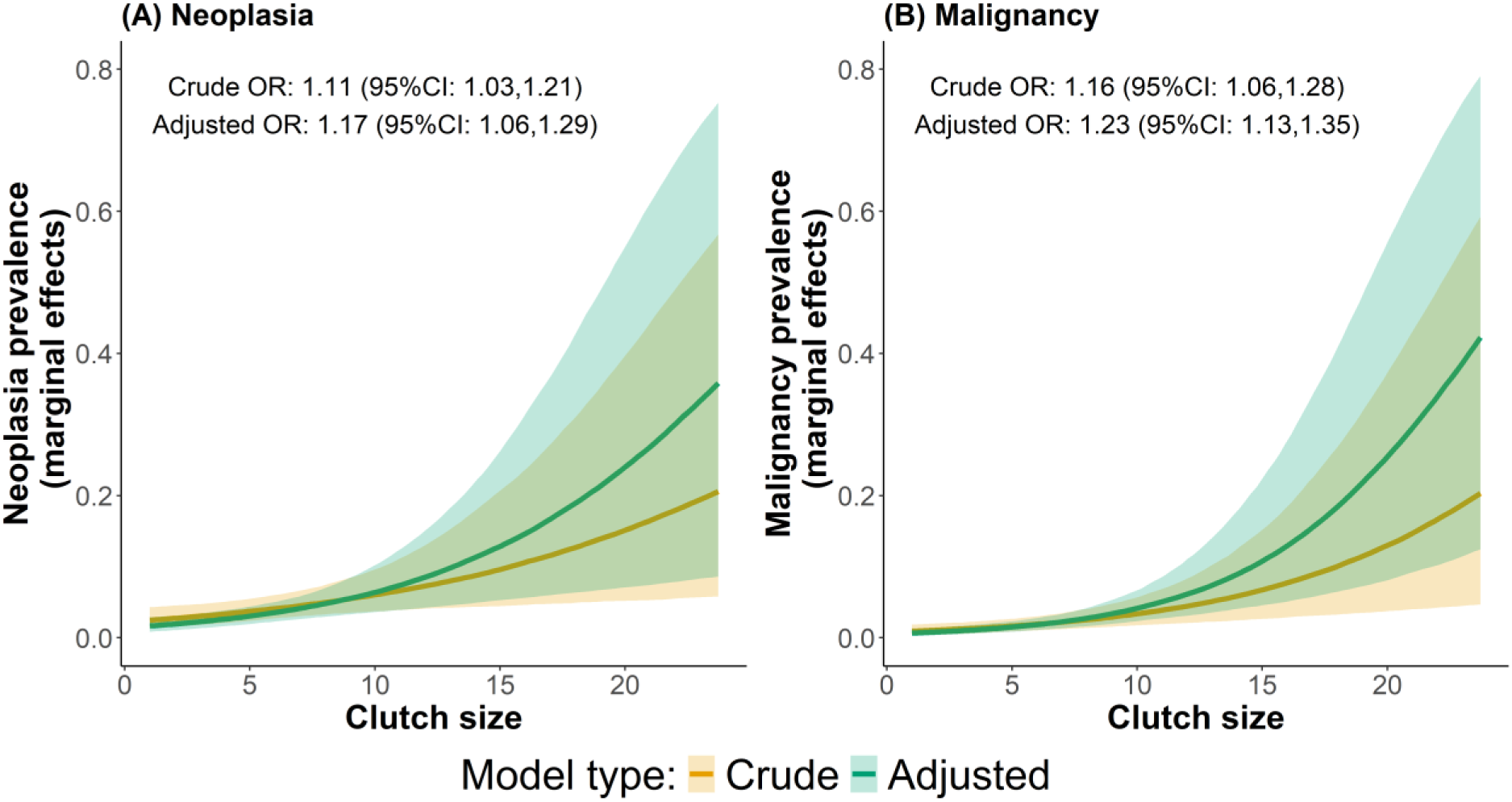
Effect of clutch size on (a) neoplasia and (b) malignancy prevalences in birds with (adjusted model) and without (crude model) accounting for confounding variables in the models. The marginals effects are showed for a species of median scientific and general public popularity.

## Discussion

Comparative oncology is a rapidly expanding field with tremendous potential, making it critically important to improve the quality of our analyses to avoid impeding progress or leading the field astray. We demonstrated that including proxies of the scientific and public popularity of a species in comparative oncology studies of captive vertebrates can greatly alter the observed trends between all three types of tumour prevalence and potential risk factors, leading to results that can sometimes be interpreted in the opposite direction to those in the original publication. For example, our study demonstrates that the bold claim by Butler *et al*. (2025) by which there is no evidence for Peto’s paradox results from their failure to account for confounding factors. We indeed found that the previously suspected small effect of body mass on cancer risk was actually due to biases related to species popularity, while the absence of trends with longevity remained unchanged, thereby providing further support for Peto’s paradox. Similarly, the complete removal of the trend between germinal cell mutation rate and cancer mortality suggests strong confounding effects, which may explain why Kapsetaki *et al*. (2024) were unsure about the biological mechanisms that would have linked those two variables as the trend was spurious.

We found that accounting for confounding factors eliminated the predicted higher mortality prevalence associated with endotheliochorial placentation. Dujon *et al*. (2024) initially predicted epitheliochorial placentation, which is the most invasive one, would be associated with the highest lethal tumour prevalences. The surprising higher cancer mortality prevalences observed in endotheliochorial placentation was hypothesised to be due to the presence of adaptations to counteract cancer cells’ ability to co-opt gene networks involved in epitheliochorial placentation to increase aggressiveness, a characteristic observed in laboratory experiments (Kshitiz *et al*. 2019). It was also proposed as a potential trade-off between reproduction and cancer risk in mammals (Dujon *et al*. 2023b). Our results suggest that cancer cells may equally co-opt these networks across all placentation types. The protective effect of gestation length on cancer risk becomes much more uncertain when confounding factors are considered. Without higher quality and larger datasets, it may be difficult to confidently conclude that species with long gestation periods have evolved defences against cancer risks associated with prolonged placental invasion during pregnancy. Very interesting, when reanalysing reproductive data in birds we observed an opposite pattern. We showed that the effect of clutch size on neoplasia and malignancy prevalence was largely underestimated, suggesting that the reproductive investment involved in egg production may be a key driver of cancer prevalence in these species. These results demonstrate how including confounding factors can substantially impact the interpretation of modelled associations between life history traits and cancer risk, not only by reducing trends providing novel information on cancer resistant, but also by revealing that some risk factors may be underestimated. Such associations have been central to discussions attempting to identify trade-offs between reproduction and cancer risk in vertebrates (Compton *et al*. 2024; Dujon *et al*. 2023b; Kapsetaki *et al*. 2024a).

Dujon *et al*. (2025) recently quantified the effect of sample size (i.e. number of necropsies) on the reliability of binomial phylogenetic models in the context of comparative oncology. We have now demonstrated that species popularity can also greatly impact those models. However, one important potential confounding factor remains unexplored and cannot be addressed with currently published datasets because prevalence estimates are pooled across all cancer types, yet cancer susceptibility is expected to vary between organs as observed in numerous species (Chu *et al*. 2012; Hubbard *et al*. 1983; Wadsworth *et al*. 1985). Organ evolution is also different across vertebrate species (Webster & Webster 2014). The patterns observed may therefore be distorted if the proportion of each cancer reported for each species is not representative of their true incidence. Future studies examining organ-specific cancer patterns will thus need to be conducted while also controlling for both scientific and public interest as confounding variables.

Unlike other disciplines that rely on cross-species comparative analyses, comparative oncology has so far overlooked some fundamental biases, an oversight that our study brings to light. The messages of this paper are therefore clear: not only do our findings substantially revise conclusions that were previously considered established, but they also emphasise that future analyses in comparative oncology should adhere to recognized best practices.

## Author contribution

AMD and FT designed the study conducted in this manuscript. AMD, JC and KA collected and consolidated the datasets. AMD conducted the analyses and wrote the initial manuscript with inputs from AMD, JC, KA and FT.

## References

1. Aktipis, A.C., Boddy, A.M., Jansen, G., Hibner, U., Hochberg, M.E., Maley, C.C., et al. (2015). Cancer across the tree of life: Cooperation and cheating in multicellularity. Philosophical Transactions of the Royal Society B: Biological Sciences, 370, 20140219.

2. Albanes, D. & Winick, M. (1988). Are cell number and cell proliferation risk factors for cancer? J Natl Cancer Inst, 80.

3. Albuquerque, T.A.F., Drummond do Val, L., Doherty, A. & de Magalhães, J.P. (2018). From humans to hydra: patterns of cancer across the tree of life. Biological Reviews, 93, 1715–1734.

4. Boddy, A.M., Abegglen, L.M., Pessier, A.P., Aktipis, A., Schiffman, J.D., Maley, C.C., et al. (2020a). Lifetime cancer prevalence and life history traits in mammals. Evol Med Public Health, 2020, 187–195.

5. Boddy, A.M., Harrison, T.M. & Abegglen, L.M. (2020b). Comparative Oncology: New Insights into an Ancient Disease. iScience, 23, 101373.

6. Boddy, A.M., Kokko, H., Breden, F., Wilkinson, G.S. & Aktipis, C.A. (2015). Cancer susceptibility and reproductive trade-offs: a model of the evolution of cancer defences. Philosophical Transactions of the Royal Society B: Biological Sciences, 370, 20140220.

7. Boutry, J., Dujon, A.M., Gerard, A.-L., Tissot, S., Macdonald, N., Schultz, A., et al. (2020). Ecological and Evolutionary Consequences of Anticancer Adaptations. iScience, 23, 101716.

8. Bürkner, P.C. (2017). brms: An R package for Bayesian multilevel models using Stan. J Stat Softw, 80, 1–28.

9. Bursac, Z., Gauss, C.H., Williams, D.K. & Hosmer, D.W. (2008). Purposeful selection of variables in logistic regression. Source Code Biol Med, 3, 1–8.

10. Butler, G., Baker, J., Amend, S.R., Pienta, K.J. & Venditti, C. (2025). No evidence for Peto’s paradox in terrestrial vertebrates. Proceedings of the National Academy of Sciences, 122, e2422861122.

11. Caulin, A.F. & Maley, C.C. (2011). Peto’s paradox: evolution’s prescription for cancer prevention. Trends Ecology Evolution, 26, 175–182.

12. Chu, P.Y., Zhuo, Y.X., Wang, F.I., Jeng, C.R., Pang, V.F., Chang, P.H., et al. (2012). Spontaneous neoplasms in zoo mammals, birds, and reptiles in Taiwan – A 10-year survey. Animal Biology, 62, 95–110.

13. Clauss, M., Roller, M., Bertelsen, M.F., Rudolf von Rohr, C., Müller, D.W.H., Schiffmann, C., et al. (2025). Zoos must embrace animal death for education and conservation. Proceedings of the National Academy of Sciences, 122, e2414565121.

14. Compton, Z.T., Mellon, W., Harris, V.K., Rupp, S., Mallo, D., Kapsetaki, S.E., et al. (2024). Cancer Prevalence across Vertebrates. Cancer Discov, OF1–OF18.

15. Compton, Z.T., Vincze, O., Mellon, W., Tollis, M., Abegglen, L., Schiffman, J.D., et al. (2025). Paradoxical indeed. Proceedings of the National Academy of Sciences, 122, e2504512122.

16. Conn, P.B., Johnson, D.S., Williams, P.J., Melin, S.R. & Hooten, M.B. (2018). A guide to Bayesian model checking for ecologists. Ecol Monogr, 88, 526–542.

17. Ducatez, S. & Lefebvre, L. (2014). Patterns of research effort in birds. PLoS One, 9, e89955.

18. Dujon, A.M., Aktipis, A., Alix-Panabières, C., Amend, S.R., Boddy, A.M., Brown, J.S., et al. (2021). Identifying key questions in the ecology and evolution of cancer. Evol Appl, 14, 877–892.

19. Dujon, A.M., Biro, P.A., Ujvari, B. & Thomas, F. (2025). Towards a more robust comparative oncology: a Bayesian reanalysis of Peto’s paradox and discussion of comparative cancer risk studies in vertebrates. R Soc Open Sci, 12, 250840.

20. Dujon, A.M., Boddy, A.M., Hamede, R., Ujvari, B. & Thomas, F. (2024a). Beyond Peto’s paradox: expanding the study of cancer resistance across species. Evolution, 79, 6– 10.

21. Dujon, A.M., Boutry, J., Tissot, S., Lemaître, J.-F., Boddy, A.M., Gérard, A.-L., et al. (2022). Cancer Susceptibility as a Cost of Reproduction and Contributor to Life History Evolution. Front Ecol Evol, 10, 861103.

22. Dujon, A.M., Jeanjean, J., Vincze, O., Giraudeau, M., Lemaître, J., Pujol, P., et al. (2023a). Cancer hygiene hypothesis: A test from wild captive mammals. Ecol Evol, 13, 1–10.

23. Dujon, A.M., Ujvari, B., Tissot, S., Meliani, J., Rieu, O., Stepanskyy, N., et al. (2024b). The complex effects of modern oncogenic environments on the fitness, evolution and conservation of wildlife species. Evol Appl, 17, e13763.

24. Dujon, A.M., Vincze, O., Lemaitre, J.-F., Alix-Panabières, C., Pujol, P., Giraudeau, M., et al. (2023b). The effect of placentation type, litter size, lactation and gestation length on cancer risk in mammals. Proceedings of the Royal Society B: Biological Sciences, 290, 20230940.

25. Hamede, R., Owen, R., Siddle, H., Peck, S., Jones, M., Dujon, A.M., et al. (2020). The ecology and evolution of wildlife cancers: applications for management and conservation. Evol Appl, 13, 1719–1732.

26. Hickisch, R., Hodgetts, T., Johnson, P.J., Sillero-Zubiri, C., Tockner, K. & Macdonald, D.W. (2019). Effects of publication bias on conservation planning. Conservation Biology, 33, 1151–1163.

27. Hirsch, J.E. (2005). An index to quantify an individual’s scientific research output. Proceedings of the National Academy of Sciences, 102, 16569–16572.

28. Hosmer Jr., D.W., Lemeshow, S. & Sturdivant, R.X. (2013). Model-Building Strategies and Methods for Logistic Regression. In: Applied Logistic Regression. John Wiley & Sons, Inc., pp. 89–151.

29. Hubbard, G.B., Schmidt, R.E. & Fletcher, K.C. (1983). Neoplasia in Zoo Animals. The Journal of Zoo Animal Medicine, 14, 33–40.

30. Hughes, A.C., Orr, M.C., Ma, K., Costello, M.J., Waller, J., Provoost, P., et al. (2021). Sampling biases shape our view of the natural world. Ecography, 44, 1259–1269.

31. Hutchins, M. (2006). Death at the zoo: The media, science, and reality. Zoo Biol, 25, 101– 115.

32. Jacqueline, C., Biro, P.A., Beckmann, C., Moller, A.P., Renaud, F., Sorci, G., et al. (2017). Cancer: A disease at the crossroads of trade-offs. Evol Appl, 10, 215–225.

33. Kamiya, T., O’Dwyer, K., Nakagawa, S. & Poulin, R. (2014). What determines species richness of parasitic organisms? A meta-analysis across animal, plant and fungal hosts. Biological Reviews, 89, 123–134.

34. Kapsetaki, S.E., Basile, A.J., Compton, Z.T., Rupp, S.M., Duke, E.G., Boddy, A.M., et al. (2025). The relationship between diet, plasma glucose, and cancer prevalence across vertebrates. Nat Commun, 16, 2271.

35. Kapsetaki, S.E., Compton, Z.T., Dolan, J., Harris, V.Κ., Mellon, W., Rupp, S.M., et al. (2024a). Life history traits and cancer prevalence in birds. Evol Med Public Health, 12, 105–116.

36. Kapsetaki, S.E., Compton, Z.T., Mellon, W., Vincze, O., Giraudeau, M., Harrison, T.M., et al. (2024b). Germline mutation rate predicts cancer mortality across 37 vertebrate species. Evol Med Public Health, 12, 122–128.

37. Kshitiz, Afzal J., Maziarz, J.D., Hamidzadeh, A., Liang, C., Erkenbrack, E.M., et al. (2019). Evolution of placental invasion and cancer metastasis are causally linked. Nat Ecol Evol, 3, 1743–1753.

38. Lynch, V.J. (2025). Peto’s paradox revisited (revisited, revisited, revisited, and revisited yet again). Proc Natl Acad Sci U S A, 122, e2502696122.

39. Manuel Vazquez, J., Pena, M.T., Muhammad, B., Kraft, M., Adams, L.B. & Lynch, V.J. (2022). Parallel evolution of reduced cancer risk and tumor suppressor duplications in Xenarthra. Elife, 11, e82558.

40. Matthews, S., Nikoonejad Fard, V., Tollis, M. & Seoighe, C. (2025). Variable Gene Copy Number in Cancer-Related Pathways Is Associated With Cancer Prevalence Across Mammals. Mol Biol Evol, 42, 1–11.

41. McClanahan, T.R. & Rankin, P.S. (2016). Geography of conservation spending, biodiversity, and culture. Conserv Biol, 30, 1089–1101.

42. McKinney, M.L. (2002). Effects of national conservation spending and amount of protected area on species threat rates. Conservation Biology, 16, 539–543.

43. Mittermeier, J.C., Correia, R., Grenyer, R., Toivonen, T. & Roll, U. (2021). Using Wikipedia to measure public interest in biodiversity and conservation. Conservation Biology, 35, 412–423.

44. Mize, T.D., Doan, L. & Long, J.S. (2019). A General Framework for Comparing Predictions and Marginal Effects across Models. Sociol Methodol, 49, 152–189.

45. Møller, A.P., Erritzøe, J. & Soler, J.J. (2017). Life history, immunity, Peto’s paradox and tumours in birds. J Evol Biol, 30, 960–967.

46. Nagy, J.D., Victor, E.M. & Cropper, J.H. (2007). Why don’t all whales have cancer? A novel hypothesis resolving Peto’s paradox. Integr Comp Biol, 47, 317–328.

47. Nery, M.F., Rennó, M., Picorelli, A. & Ramos, E. (2022). A phylogenetic review of cancer resistance highlights evolutionary solutions to Peto’s Paradox. Genet Mol Biol, 45, e20220133.

48. Noble, R., Kaltz, O. & Hochberg, M.E. (2015). Peto’s paradox and human cancers. Philosophical Transactions of the Royal Society B: Biological Sciences, 370, 20150104.

49. Nunney, L. (1999). Lineage selection and the evolution of multistage carcinogenesis. Proc R Soc Lond B Biol Sci, 266, 493–498.

50. Nunney, L. (2018). Size matters: height, cell number and a person’s risk of cancer. Proceedings of the Royal Society B: Biological Sciences, 285, 20181743.

51. Nunney, L. (2022). Cancer suppression and the evolution of multiple retrogene copies of TP53 in elephants: A re-evaluation. Evol Appl, 15, 891–901.

52. Nunney, L. (2024). The effect of body size and inbreeding on cancer mortality in breeds of the domestic dog: a test of the multi-stage model of carcinogenesis. R Soc Open Sci, 11, 231356.

53. Poulin, R., Presswell, B., Bennett, J., de Angeli Dutra, D. & Salloum, P.M. (2023). Biases in parasite biodiversity research: why some helminth species attract more research than others. Int J Parasitol Parasites Wildl, 21, 89–98.

54. Roche, B., Hochberg, M.E., Caulin, A.F., Maley, C.C., Gatenby, R.A., Misse, D., et al. (2012). Natural resistance to cancers: a Darwinian hypothesis to explain Peto’s paradox. BMC Cancer, 12, 387.

55. Roll, U., Mittermeier, J.C., Diaz, G.I., Novosolov, M., Feldman, A., Itescu, Y., et al. (2016). Using Wikipedia page views to explore the cultural importance of global reptiles. Biol Conserv, 204, 42–50.

56. Seluanov, A., Gladyshev, V.N., Vijg, J. & Gorbunova, V. (2018). Mechanisms of cancer resistance in long-lived mammals. Nat Rev Cancer, 18, 433–441.

57. Simons, M.J.P. (2025). Peto’s paradox’s relevance is off the scale. Aging, 13, 357–365.

58. Szumilas, M. (2010). Explaining odds ratios. Journal of the Canadian Academy of Child and Adolescent Psychiatry, 19, 227–229.

59. Tam, J., Lagisz, M., Cornwell, W. & Nakagawa, S. (2022). Quantifying research interests in 7,521 mammalian species with h-index: a case study. Gigascience, 11, 1–11.

60. Thomas, F., Jacqueline, C., Tissot, T., Henard, M., Blanchet, S., Loot, G., et al. (2017). The importance of cancer cells for animal evolutionary ecology. Nat Ecol Evol, 1, 1592–1595.

61. Tollis, M., Boddy, A.M. & Maley, C.C. (2017). Peto’s Paradox: how has evolution solved the problem of cancer prevention? BMC Biol, 15, 1–5.

62. Trivedi, D.D., Dalai, S.K. & Bakshi, S.R. (2023). The Mystery of Cancer Resistance: A Revelation Within Nature. J Mol Evol, 91, 133–155.

63. Vincze, O., Colchero, F., Lemaître, J.-F., Conde, D.A., Pavard, S., Bieuville, M., et al. (2022). Cancer risk across mammals. Nature, 601, 263–267.

64. Wadsworth, P.F., Jones, D.M. & Pugsley, S.L. (1985). A Survey of Mammalian and Avian Neoplasms at the Zoological Society of London. The Journal of Zoo Animal Medicine, 16, 73–80.

65. Webster, Douglas. & Webster, Molly. (2014). Comparative Vertebrate Morphology. Elsevier Science.

66. Welch, H.G. & Fisher, E.S. (2017). Income and Cancer Overdiagnosis — When Too Much Care Is Harmful. New England Journal of Medicine, 376, 2208–2209.

