## Supplementary Figures 1-6 for "The Cost of Fame: Strong Biases in Comparative Oncology of Captive Species"

### Supplementary Material

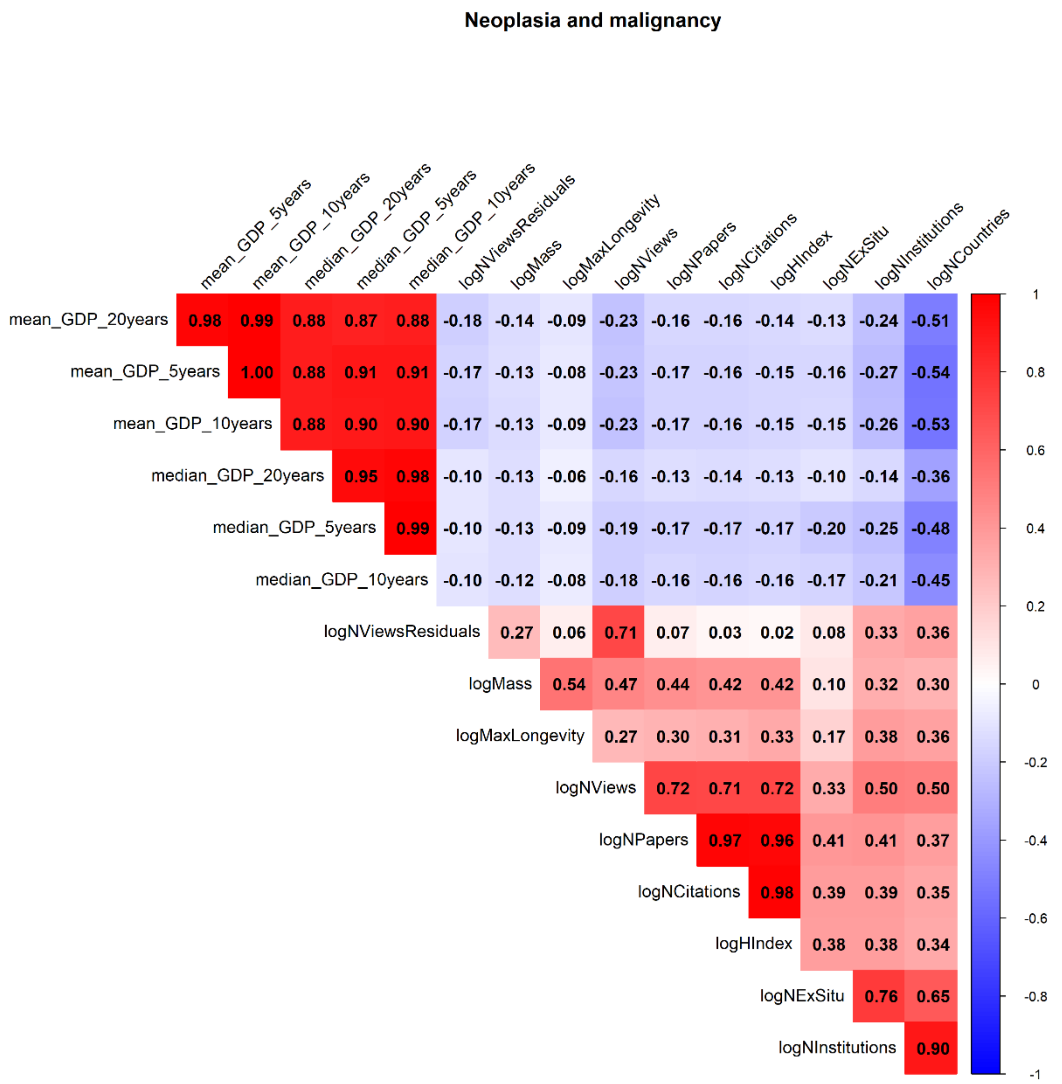


**Supplementary Figure 1:** Correlation matrix of potential confounding variables associated with neoplasia and malignancy prevalence in vertebrates.

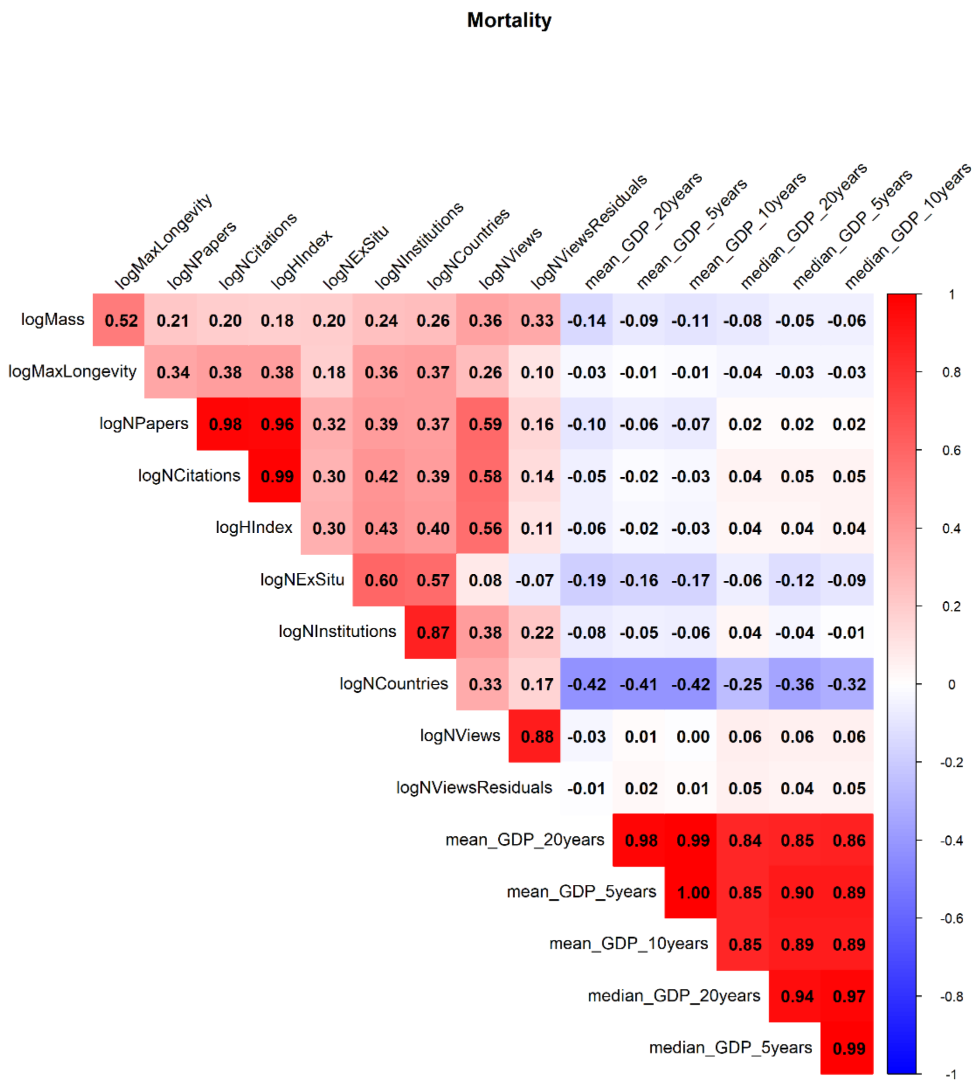


**Supplementary Figure 2:** Correlation matrix of potential confounding variables associated with cancer mortality prevalence in vertebrates.

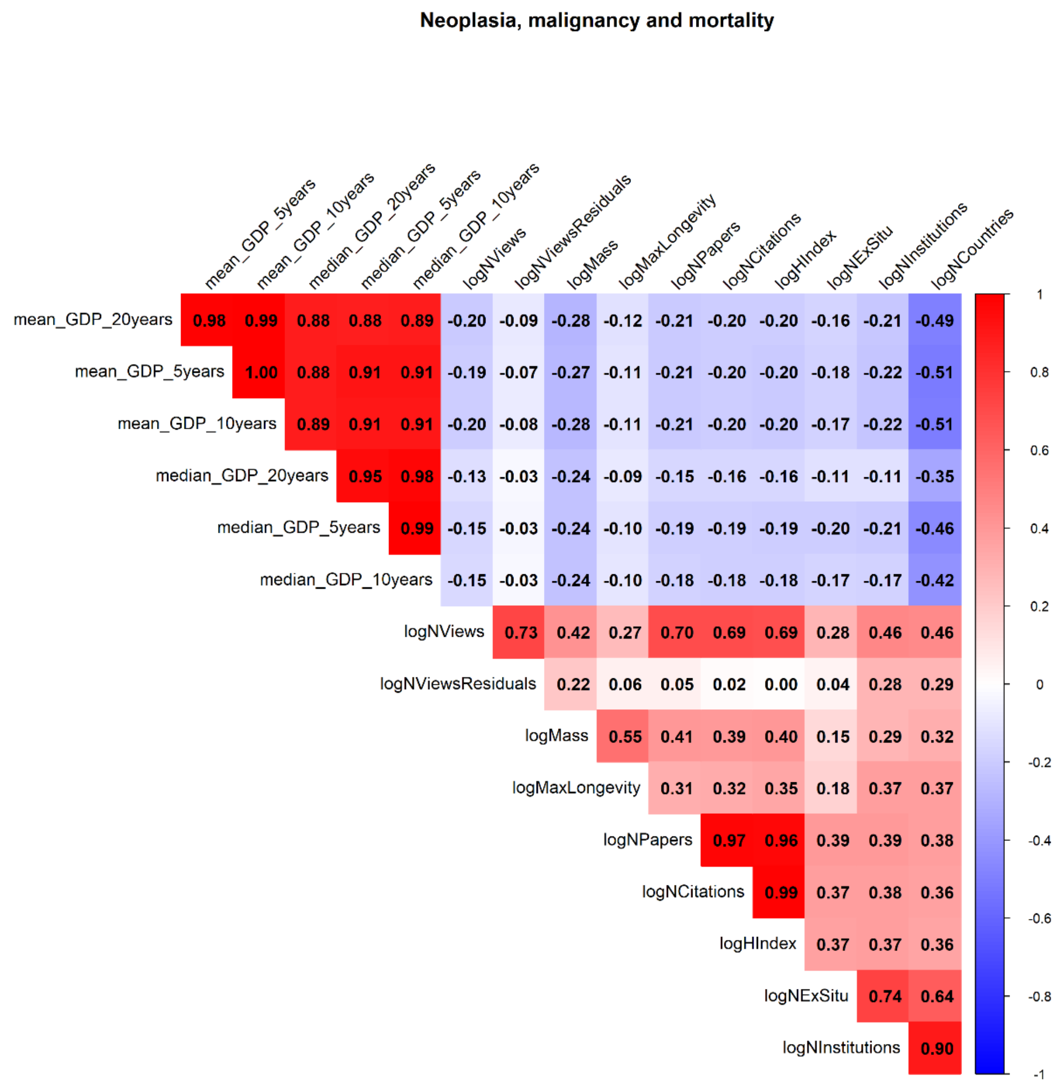


**Supplementary Figure 3:** Correlation matrix of potential confounding variables associated with neoplasia, malignancy and cancer mortality prevalence in vertebrates.

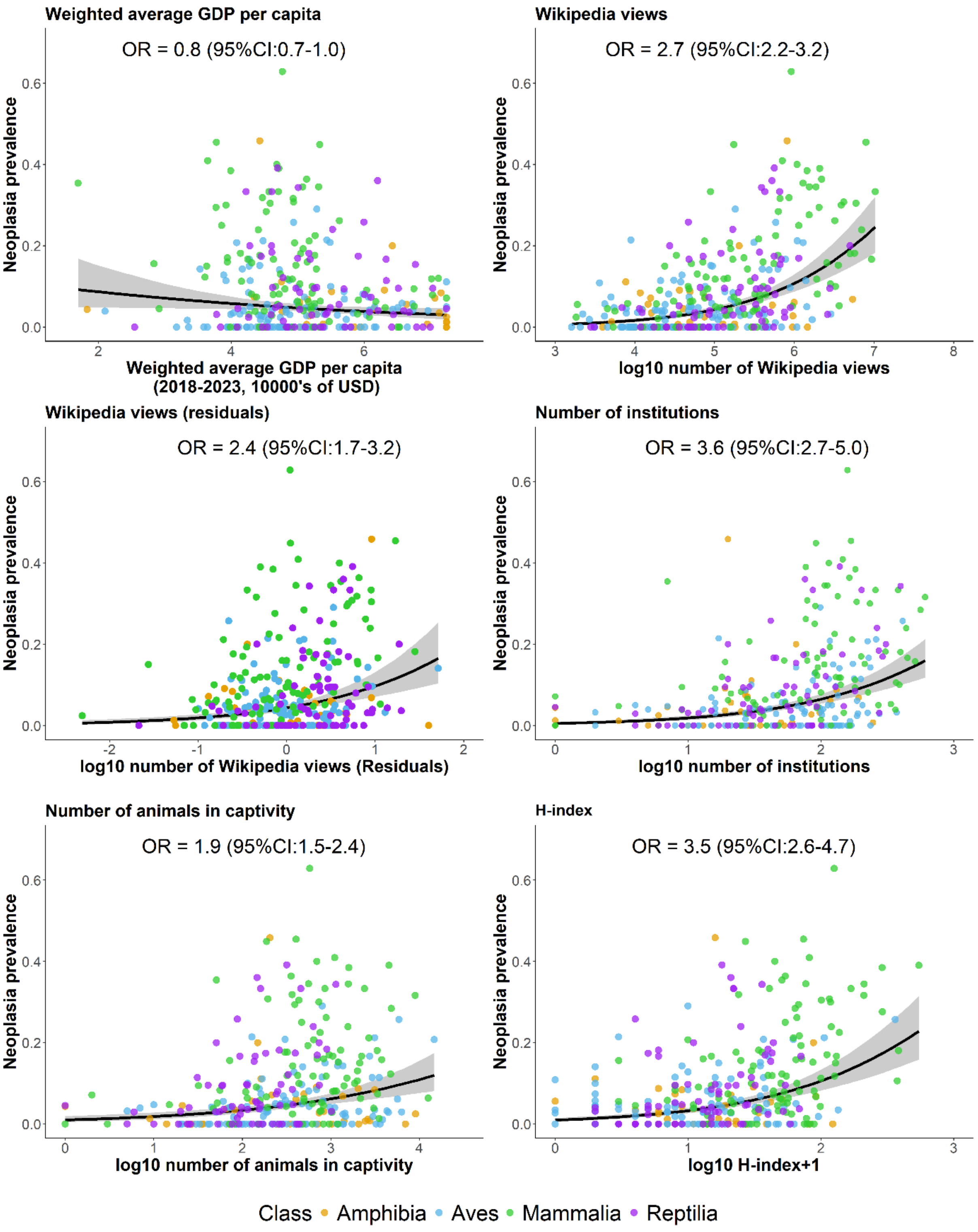


**Supplementary Figure 4:** Binomial phylogenetic regression models with 95% credible intervals bands examining the relationship between neoplasia prevalence and various proxies of scientific and public interest in captive vertebrates.

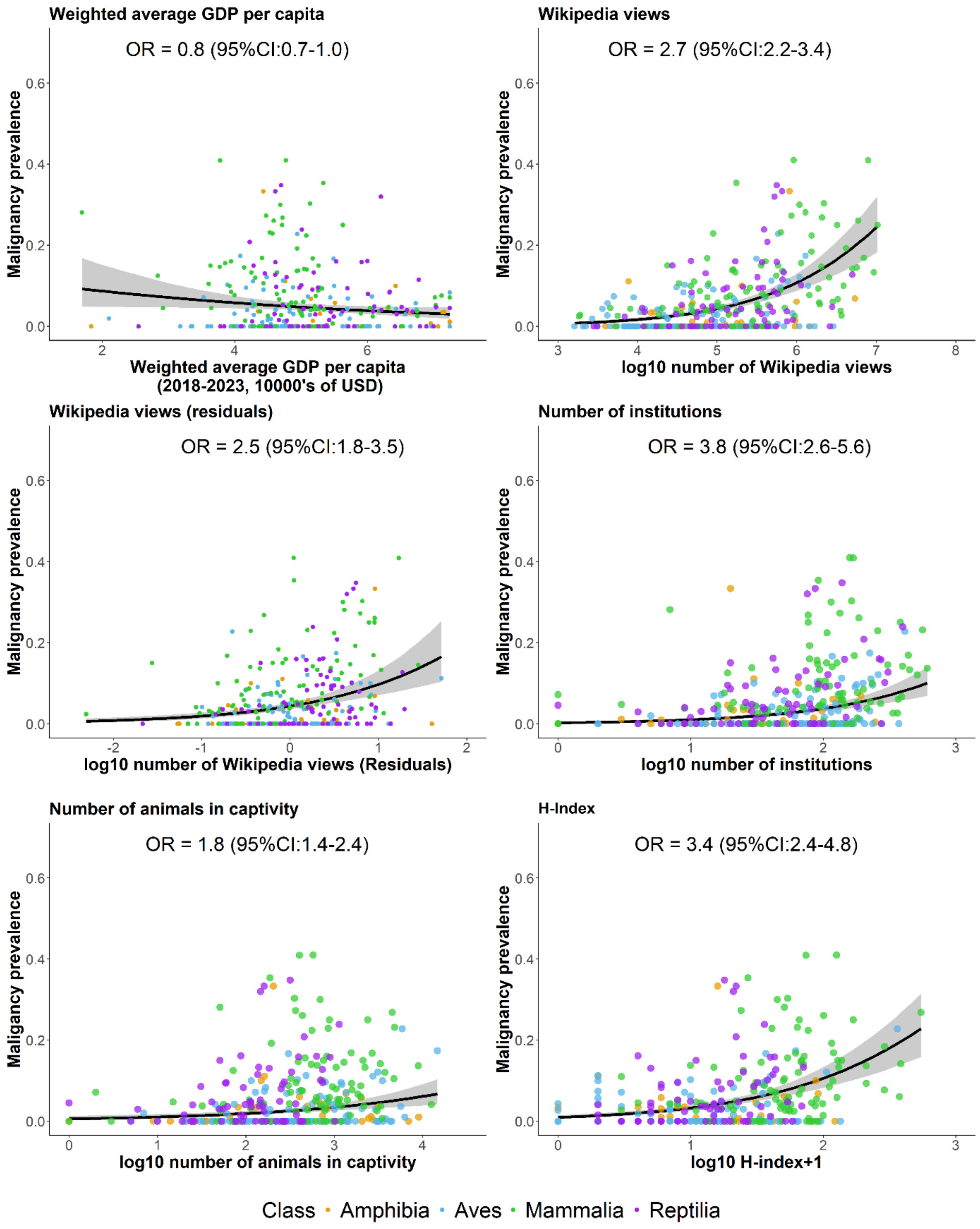


**Supplementary Figure 5:** Binomial phylogenetic regression models with 95% credible intervals bands examining the relationship between malignancy prevalence and various proxies of scientific and public interest in captive vertebrates.

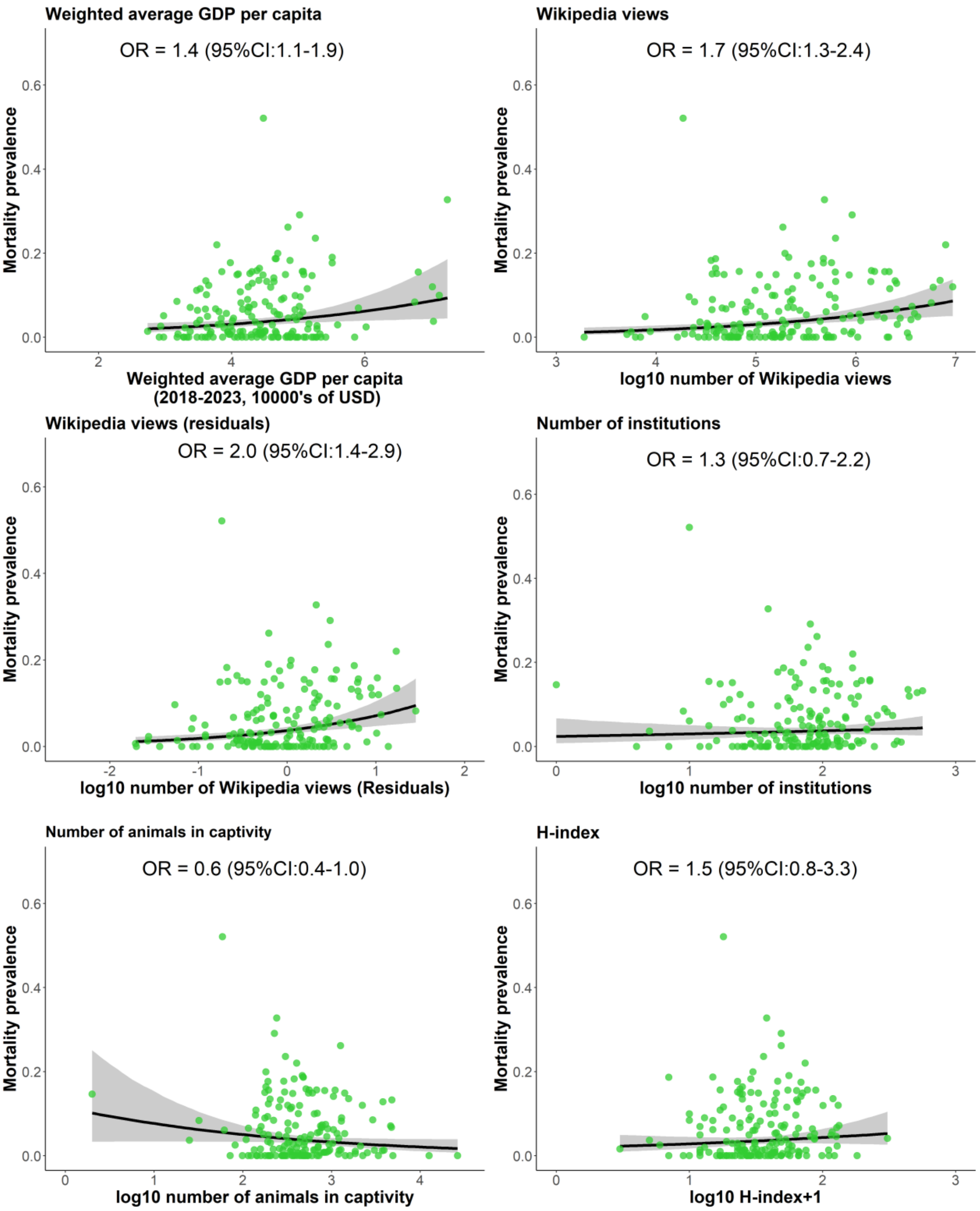


**Supplementary Figure 6:** Binomial phylogenetic regression models with 95% credible intervals bands examining the relationship between cancer mortality prevalence and various proxies of scientific and public interest in captive mammals.
